# What makes *Candida auris* pan-drug resistant? Integrative insights from genomic, transcriptomic, and phenomic analysis of clinical strains resistant to all four major classes of antifungal drugs

**DOI:** 10.1101/2024.06.18.599548

**Authors:** Johanna Rhodes, Jonathan Jacobs, Emily K. Dennis, Swati R. Manjari, Nilesh Banavali, Robert Marlow, Mohammed Anower Rokebul, Sudha Chaturvedi, Vishnu Chaturvedi

**Author notes:** Corresponding author: < >. Equal contributions (Johanna Rhodes, Jonathan Jacobs).

## Abstract

The global epidemic of drug-resistant *Candida auris* continues unabated. We do not know what caused the unprecedented appearance of pan-drug resistant (PDR) *Candida auris* strains in a hospitalized patient in New York; the initial report highlighted both known and unique mutations in the prominent gene targets of azoles, amphotericin B, echinocandins, and flucytosine antifungal drugs. However, the factors that allow *C. auris* to acquire multi-drug resistance and pan-drug resistance are not known. Therefore, we conducted a comprehensive genomic, transcriptomic, and phenomic analysis to better understand PDR *C. auris*. Among 1,570 genetic variants in drug-resistant *C. auris*, 299 were unique to PDR strains. The whole genome sequencing results suggested perturbations in genes associated with nucleotide biosynthesis, mRNA processing, and nuclear export of mRNA. Whole transcriptome sequencing of PDR *C. auris* revealed two genes to be significantly differentially expressed - a DNA repair protein and DNA replication-dependent chromatin assembly factor 1. Of 59 novel transcripts, 12 candidate transcripts had no known homology among expressed transcripts found in other organisms. We observed no fitness defects among multi-drug resistant (MDR) and PDR *C. auris* strains grown in nutrient-deficient or - enriched media at different temperatures. Phenotypic profiling revealed wider adaptability to nitrogenous nutrients with an uptick in the utilization of substrates critical in upper glycolysis and tricarboxylic acid cycle. Structural modelling of 33-amino acid deletion in the gene for uracil phosphoribosyl transferase suggested an alternate route in *C. auris* to generate uracil monophosphate that does not accommodate 5-fluorouracil as a substrate. Overall, we find evidence of metabolic adaptations in MDR and PDR *C. auris* in response to antifungal drug lethality without deleterious fitness costs.

## Introduction

*Candida auris* became a global threat agent in the decade since its 2009 discovery from the ear canal of a Japanese patient (1–3). It’s rapid geographic spread, ability to persist in the hospital environments, and acquisition of multi-drug resistance distinguishes *C. auris* from all other known human fungal pathogens (4–8). In the United States, the emergence and transmission of multi-drug resistant (MDR) *C. auris* strains, which were non-responsive to most commonly used azole and echinocandin drugs have been a cause for concern (9–11). Our most recent report documented an unprecedented resistance to the four most commonly available classes of antifungal drugs in *C. auris* pan-drug resistant (PDR) strains from a transplant patient in New York (12).

Current emphasis on the control and management of *C. auris* in healthcare facilities rely on rapid diagnosis by molecular methods, environmental decontamination, and the judicious use of antifungal drugs (13–16). Molecular tests are also being developed for the rapid detection of *C. auris* drug resistance markers for azole, echinocandins, and flucytosine (5-flucytosine, 5-FC) (12, 17–19). However, despite these efforts, the continuing spread of drug resistant *C. auris* in the US and other parts of the world demands immediate attention (20, 21). There remain several gaps in the understanding of *C. auris* disease processes and filling them could improve its control and treatment options.

Several investigators have reported experimental studies with drug-resistant *C. auris* to assign critical roles for drug target genes *ERG11* and *FKS1*, efflux pumps activations, biofilms, differential gene expression and structural alterations, lipid perturbations, aneuploidy, and chromatin modifications (7, 22–27). A similar approach is not yet published with clinical *C. auris* strains resistant to all four classes of widely available antifungal drugs. Therefore, we hypothesized that *C. auris* strains from our patient could provide insights into the genomic, transcriptomic, and phenomic signatures driving the acquisition of pan-drug resistance.

## Methods

### Candida auris

Two *C. auris* strains, selected for this study, originated from a transplant patient described in an earlier publication by Jacobs *et al*. (12). The initial *C. auris* strain20-34 (MDR), was collected four days after hospitalization and, the strain *C. auris* 20-32 (PDR) was collected 69 days after hospitalization. Both strains were confirmed as *C. auris* Clade I by Sanger sequencing of ITS and D1/D2 ribosomal genes, and MALDI-TOF-MS (28). Breakpoints for fluconazole, echinocandins, and amphotericin B were interpreted per CDC recommendations. Antifungal susceptibility testing was carried out as described earlier by Jacobs *et al.* (12).

### Fungal growth

Two *C. auris* strains were tested for fitness by monitoring growth in RPMI1640 medium and Sabouraud broth at 30 °C and 37 °C. For biosafety reasons, we preferred stationary cultures in 96-well microtiter plates (9). The plates were read without opening lids at periodic intervals with a BioTek Synergy 2 plate reader to record _A750._ Data analysis and plotting was done with the GraphPad Prism 8 software for Mac.

### Whole genome sequencing and bioinformatic analysis

Whole genome sequencing and variant calling of *C. auris* 20-34 and 20-32 was performed using the *C. auris* B8441v2 reference genome (PEKT02000000)(5). Whole genome sequencing was performed for each strain as described earlier in Jacobs *et al.* (12).

### RNAseq library preparation

Briefly, *C. auris* 20-34 and 20-32 were streaked on Sabouraud dextrose agar and incubated for 24□h at 35°C. The yeast cells were suspended in sterile water to an absorbance (A_530_) of 0.08 to 0.1 as measured with a Mettler Toledo UV5 bio spectrophotometer; 20□µl of the cell suspension was added to 11□ml of RPMI 1640 broth tube. One hundred microliters of the suspension in RPMI 1640 was added to each well of the 96-well plate containing 100 µl RPMI 1640 and incubated in triplicate for 0 hrs and 6 hrs. At the end of the incubation, the cell suspension from each well was harvested by centrifugation at 12,500 rpm for 1 min. Total RNA extractions were done using Epicenter MasterPure Yeast RNA Purification Kit (Lucigen, Middleton, WI). The RNA samples were treated with DNase I to remove all traces of DNA using TURBO DNA-free™ Kit (Invitrogen™, Carlsbad, CA) in accordance with the manufacturer’s instructions. The RNA samples were purified for RNA sequencing to be free of salts (e.g., Mg2+, or guanidinium salts), divalent cation chelating agents (e.g., EDTA, EGTA, citrate), or organics (e.g., phenol and ethanol). Total purified RNA was quantified and quality checks (QCs) using NanoDrop™ 2000/2000c Spectrophotometers (Thermo Scientific™, Waltham, MA) and Qubit™ RNA HS Assay Kit (Thermo Scientific™, Waltham, MA) as manufacturer’s instructions. Further to assure the RNA quality and integrity, the RNA samples were run on an Agilent Bioanalyzer→ RNA 6000 Nano/Pico Chip and the final concentration of purified RNA were measured.

We also exposed *C. auris* 20-34 and 20-32 to 5-FC at final concentration of 2 µg/mL, which represents the high minimum inhibitory concentration (MIC) of the drug, and separates flucytosine-susceptible strain (*C. auris* 20-34) from the -flucytosine-resistant strain (*C. auris* 20-32) (12). The cells were processed in triplicate as described earlier for non-drug exposure treatment.

Libraries for whole transcriptome sequencing were produced from 20 ng total RNA per sample using NEBNext Ultra II RNA Library Prep Kit for Illumina (New England Biolabs) and its corresponding sample indexing kit (Index Primer Set 1). Finished libraries were assessed for quality prior to sequencing (a) *via* quantitative analysis using Qubit Broad Range assay kit, and (b) for library preparation size distribution *via* Agilent 4200 TapeStation. Libraries were then sequenced on the Illumina NextSeq 2000 platform using paired-end 2 x 150 bp sequencing reads with Illumina Stranded mRNA Prep kit. The raw fastq data have been deposited under NCBI BIOPROJECT ID PRJEB57846.

### Whole transcriptome Ranse analysis

The splice-aware aligner HISAT2 v2.2.1 (29) was used to align sequencing reads to the *C. auris* B8441 (5) reference genome (GCA_002759435.2). Annotated gene abundances were quantified using StringTie v1.3.3b (30), and annotated reference genes were identified using the *C. auris* B8441 GFF3 annotation file.

Gene expression levels were quantified using Ballgown v3.15 (20) in R v4.2.1 based on Fragments Per Kilobase Million (FPKM). Low abundance genes, classed as genes with transcripts with a variance across the samples of less than one, were filtered out. Expression levels were log transformed, and significant differentially expressed genes with a *p*-value < 0.05 were retained.

Overrepresented gene ontology (GO) annotation amongst up and down-regulated genes in each condition were predicted using BiNGO (using a custom GO Full annotation for *C. auris* available at: https://doi.org/10.5281/zenodo.10137315) in Cytoscape v3.9.1 using hypergeometric exact tests with an FDR ≤ 0.05. Protein domains were identified using HMMER v3.3.2 (31) with default settings against the Pfam database (32).

### Novel transcript discovery

Novel transcript discovery was carried out for both *C. auris* 20-32 and 20-34, which resulted in the identification of candidate novel transcripts for a detailed future study. Briefly, the raw FASTQ sequencing data for RNAseq (see above) was combined for each strain across each condition and were imported into CLC Genomics Server v23 (QIAGEN Digital Insights, https://resources.qiagenbioinformatics.com/manuals/transcriptdiscovery/current/index.php) as a two data sets (one for each strain). A total of 516,619,984 and 504,865,706 paired-end reads were used for each strain. Combined raw reads were then trimmed and filtered in CLC using the default settings. Trimmed reads were subsequently mapped to *C. auris* B8441 (5)reference genome (GCA_002759435.2) using CLC’s Large Gap Read Mapping tool with the default settings and a distance max of 50 kb. Following mapping, CLC’s Transcript Discovery Tool was run on each strain’s mapping data using the following settings: Minimum ORF length, 100; Ignore Duplicates; Ignore non-specific matches; Minimum Spliced Reads, 1; Minimum Spliced Coverage, 0.05; Ignore chimeric reads; Minimum Unspliced Reads, 5; Minimum Unspliced Coverage, 0.05; Gene merging distance, 50; Minimum reads, 10; Minimum predicted gene length, 250; Ignore genes with no spliced reads; Exon merging distance = 100. Next, CLC’s Detect & Refine Gene Fusion Tool was run on each of the input datasets with the following settings: Max fusions per event, 100; Minimum unaligned read count, 1; Minimum unaligned read length, 15; Maximum distance for broken pair fusions, 1000; Assumed Error Rate, 0.001; Promiscuity Threshold, 7; Detect Exon Skipping; Detect Novel Exon Boundaries; Minimum number of supporting reads per fusion, 2; Max p-value, 0.05; Min. Z-score, 2.5; Breakpoint Distance, 10; Skip insignificant breakpoints; Mapping Cost, 2; Insertion Cost, 3; Deletion Cost, 3; Length Fraction, 0.8; Similarity Fraction, 0.8. The results of both Transcript Discovery Tool and the Detect & Refine Gene Fusion Tool were a set of candidate tracks from each strain. These tracks were then merged and a manually reviewed. Candidate transcripts that were found to overlap or were predicted to be a potential sub-genomic promoter start sites were noted and removed from further consideration. Following manual review, all candidate transcripts were used as a BLAST query against both *nr* and *refseq_rna* databases at NCBI (33). Differential gene expression analysis was carried out in CLC to compare differences in potential novel transcripts expressed between strains, or in response to drug treatment, using parameters as described above.

### Whole genome variant calling

Whole genome sequencing of 20-32 and 20-34 was carried out as described in Jacobs *et al*. (12). Variant calling for each strain was done using B8441v2 as a common reference (5). Briefly, 2x150 bp paired-end Illumina reads were trimmed and filtered to Q30 as previously described. Reads from 20-32 and 20-34 libraries where then mapped to the B8441v2 reference genome to an average coverage depth of 50x and 171x coverage respectively. Indel and Structural Variant analysis was done (*p*-value < 0.0001) with B8441v2 used as a guidance track. To improve final variant calling confidence, reads mapped to all candidate variants, indels, and structural variations were iteratively locally realigned with a minimum alignment window of 200 bp. Variant detection was then carried out with the following parameters: Ignore broken and non-specific paired reads; coverage > 10; count > 2,; frequency > 66.%, quality >= Q30 within 30 nt window; bidirectional reads >5%, read balance direction p(x)< 0.01; read start position balance p(x)< 0.01. All variant calling was carried out using CLC Biomedical Genomics Tools (QIAGEN Digital Insights, Aarhus, Denmark).

### Phenotypic array

We utilized Biolog PM microarray plates to confirm *in silico* gene predictions per an earlier publication from our laboratory (34). Two *C. auris* strains, 20-34 and 20-32, were grown on SAB agar at 37 °C. A cell suspension for each *C. auris* strain was made in sterile water and adjusted to a transmittance of 62% (OD_600_). Biolog Phenotypic Microarray plates (Biolog Inc., Hayward, CA) were inoculated and incubated at 37 °C in a stationary incubator and A_490_ recorded (BioTek Synergy 2, Winooski, VT, USA). End point readings for carbon sources (PM1 and PM2 plates) and nitrogen sources (PM3) were collected after 72 h. The average negative control value for that timepoint was subtracted from the well absorbances (34). Two trials for PM1 on two separate days were averaged and those values are presented. Only one replicate was run for PM2 and PM3.

### Modelling

A structural model of the tetramer for *C. auris* uracil phosphoribosyl transferase (UPRT) was constructed using homology modelling through Swiss-Model workspace ((35), https://swissmodel.expasy.org) with the *Toxoplasma gondi* (*Tgo*) UPRT crystal structure, PDB ID: 1JLS (36), as a template. The *Tgo* UPRT structure, with a resolution of 2.5 Å, was chosen over the unpublished *Candida albicans* (*Cal*) UPRT structure (PDB ID: 7RH8), with a resolution of 1.7 Å, because a *Tgo* UPRT monomer has two bound substrates, uracil and phosphoribosyl pyrophosphate (PRPP), which represents a better reactant state model. Models of the *C. auris* UPRT are available (Uniprot IDs: A0A2H0ZP91, A0A0L0P8H5, A0A510NX92) in the AlphaFold2 database (37, 38)(https://alphafold.ebi.ac.uk), but these represent monomer models rather than tetramer models, and also do not take into account any adjustments to bound substrates. The substrate ligand structures in the *C. auris* UPRT tetramer model were obtained by aligning the individual *C. auris* UPRT monomers to their *Tgo* UPRT monomer counterparts by using the matchmaker module in ChimeraX v1.4 (39). The ligand models were built in Phenix v1.16 (40) using the phenix.elbow module (41) and the entire tetramer was minimized for 20 macrocycles without any restraints to get the final tetramer model.

## Results

### Genomics of pandrug-resistant *C. auris* reveals mutations in genes involved in antifungal drug resistance

The number of single nucleotide polymorphisms (SNPs), insertions and deletions (indels) and structural variants (SVs) found in the Day four (*C. auris* 20-34, MDR) and Day 69 (*C. auris* 20-30, *C. auris* 20-31 and *C. auris* 20-32, all PDR) strains, are shown in Table 1. Collectively, all four strains shared 1570 variants compared to the reference strain. All three Day 69 strains collectively had 299 variants among them that differed from the Day four strain, 82 variants were common to all Day 69 strains, and were the focus of further analysis (Figure 1A, Supp Data 1).

**Figure 1:**
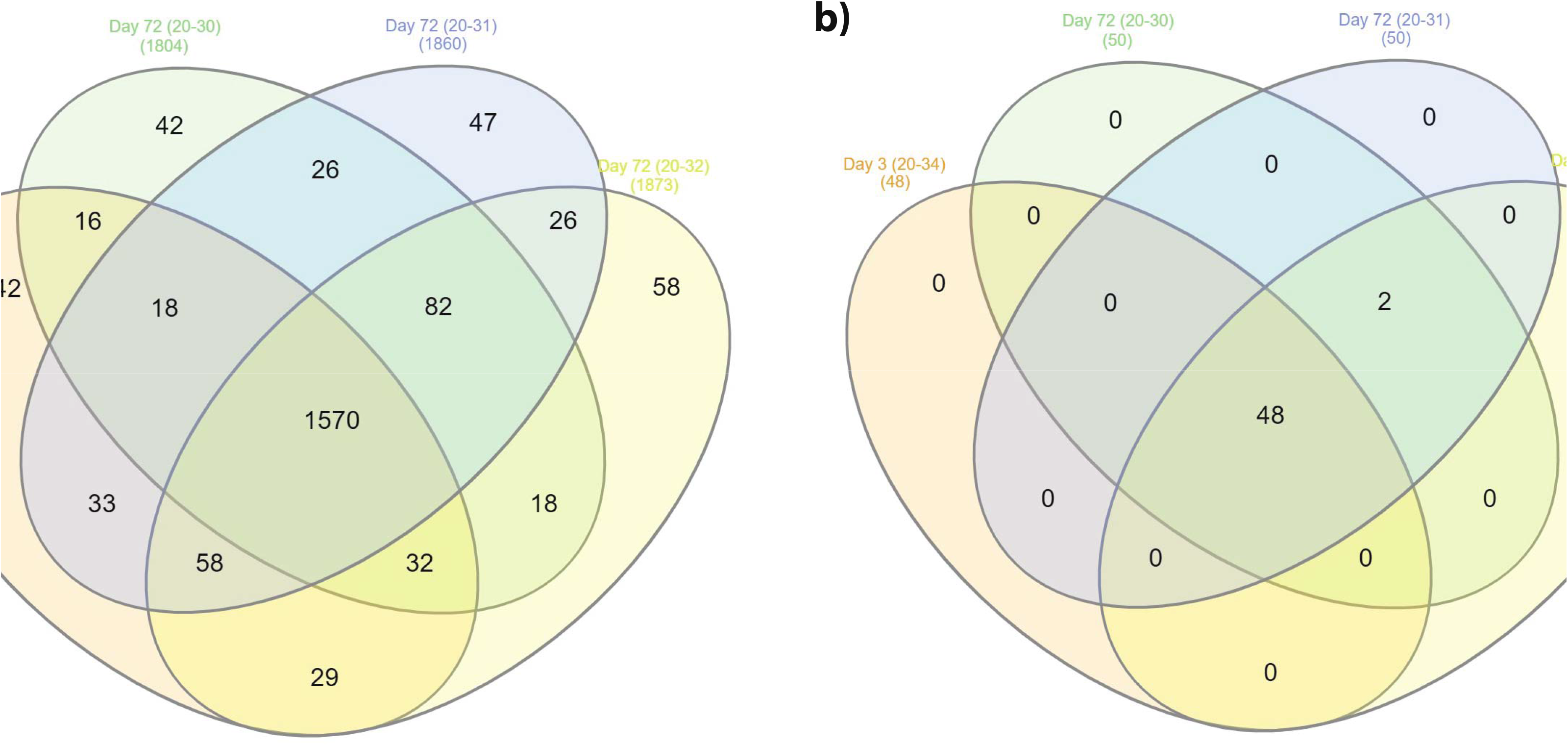
Whole genome analysis of *C. auris* strains 20-30, 20-31, 20-32 and 20-34. A) shared variants (SNPs, indels and SVs) between Day 4 (20–34) and Day 69 (20-30, 20-31 and 20-32) strains B) Gene set enrichment analysis (GSEA) for genes with variants for Day 4 and Day 69 strains.

**Table 1.** Variants (SNPs, indels and SVs) in Day 4 (*C. auris* 20-34) and Day 69 (*C. auris* 20-30, 20-31 and 20-32) strains.

|  | *Candida auris* |  |  |  | Shared<br>ALL | Unique<br>to Day<br>69 | Shared<br>Among<br>Day 69<br>Only |
| --- | --- | --- | --- | --- | --- | --- | --- |
|  | 20-30 | 20-31 | 20-32 | 20-34 |  |  |  |
| SNV | 1,030 | 1,044 | 1,046 | 1,045 | 989 | 57 | 16 |
| MNV | 18 | 18 | 19 | 18 | 14 | 6 | 1 |
| Insertion | 183 | 196 | 194 | 166 | 145 | 53 | 23 |
| Deletion | 569 | 598 | 609 | 666 | 419 | 178 | 41 |
| Replacement | 4 | 4 | 5 | 3 | 3 | 1 | 1 |
| Total (non-reference) | 1,804 | 1,860 | 1,873 | 1,898 | 1,570 | 295 | 82 |

Gene set enrichment analysis (GSEA) for genes with variants revealed two enriched gene ontology (GO) terms specific to Day 69, and an additional 48 GO terms enriched collectively among the four strains (*p*-value < 0.05, Figure 1B). Only two terms were shared for the Day 69 strains: pyrimidine-containing compound salvage pathway (GO:0008655) and poly(A)+ mRNA export from nucleus (GO:0016973). These results correspond to mutations in B9J08_004076, B9J08_001846, and B9J08_003760 that are shared among all three Day 69 strains but absent from the Day 4 strain (Figure 1B).

All three Day 69 strains exhibits the same deletion of a 131 bp region in the 5’ end of B9J08_004076, encoding *FUR1*, the uracil phosphoribosyltransferase (UPRT) catalysing the first step in UMP biosynthesis, and the target of the antifungal drug flucytosine. This deletion also included the promoter region (Supp Fig 1A), and was predicted to totally disrupt expression and function of *FUR1* in these strains, likely significantly contributing to its resistance to nucleoside analogues, including flucytosine.

There was also a common mutation (PIS55741.1:c.1579dupC) in all three Day 69 strains that is absent in the Day four strain (Supp Fig 1B), which would result in a frameshift mutation (Ser527fs) near the terminus of the protein, potentially impacting function. BLAST analysis of PIS55741.1 revealed similarity to both a putative mRNA export factor in *Clavispora lustianiae* (OVF10997.1, 57% identity, 72% aligned, E-value 0.00) and the nuclear export factor *Mex67* in *Candida orthopsilosis* (XP_003869218.1, 50% identity, 65% aligned, E-value 2e^-172^). The impact of a mutation in the terminal domain of *Mex67* may disrupt its ability to bind nuclear pore proteins (i.e. nucleoporins) and the shuttling of mRNAs out of the nucleus.

The third mutation present in all Day 69 strains was a 15 nt in-frame deletion in B9J08_003760, predicted to result in PIS52149.1:p.Gln444_Gly448del. The impact of this mutation is not known, but analysis using CDART (42) suggests that the encoded protein PIS52149.1 functions as a nucleoporin, again highlighting that defects in mRNA export from the nucleus may be playing a role in resistance to flucytosine and other antifungal drugs.

In comparison to the *C. auris* B8441v2 reference genome, 20-32 and 20-34 had 1,804 and 1,898 variants respectively, with 1,570 variants shared between them (Table 1). The 20-32 PDR strain was found to have 82 variants not found in 20-34, including 66 in or adjacent to homopolymer regions and 3 tandem repeats which were not considered for further analysis. The remaining mutations included five non-synonymous mutations, including the previously reported 139 bp deletion in the promoter region of *FUR1* (PIS52459.1: p.M1fs), as well as a T>G mutation in 1,3-beta-glucan synthase component *FKS1* (B9J08_000964, PIS58465.1: p.F635C), a G>T in bifunctional purine biosynthesis protein ADE17 (B9J08_001850, PIS55745.1: p.G45V), a 180 bp deletion in B9J08_001808 (PIS55703.1: p.A23_G82del), a 24 bp deletion in B9J08_002593 (PIS51023.1: p.I16_G23del), a T>G mutation in B9J08_000964 (PIS58465.1: p.F635C), and a 21 bp deletion in a repeat region of the hypothetical protein B9J08_004455. Interestingly, the hypothetical protein B9J08_004100 was found to have 5 mutations with sub-allelic variant frequencies ranging as low as 75%, suggesting perhaps that active selective pressure was still driving the progression of mutations in this strain. While the functional consequences of the deletion in *FUR1* and the mutations in *FKS1* and *ADE17* are expected to be significant contributing factors to the pan-resistance phenotype observed in *C. auris* 20-32, the impact of the remaining nonsynonymous mutations remains to be determined.

### Novel transcript discovery reveals significantly differentially expressed genes involved in adaptive stress response

The reference genome annotations for *C. auris* are based on strain B8441v2 (4,27). Nonetheless, due to the rapid accumulation of mutations during the course of prolonged hospitalization of the patient, we sought to identify potentially novel transcripts or splice-site isoforms that may not be present in the canonical B8441v2 reference genome. To this end, we used an established pipeline for RNAseq transcript discovery and applied this pipeline to both strains. This resulted in the identification of 59 novel transcripts not found in the B8441v2 reference genome annotation. Of these 59 transcripts, 47 were found to have some level of homology to “known transcripts” found in other organisms based on pairwise sequence identity (ranging from 67.5% to 100.0%), but were not annotated in B8441v2 genome reference. Collectively, using BLAST (23) 47 transcripts were identified as potentially being full or partial mRNAs (41), snoRNAs (5), or a pre-ribosomal miscRNA (1). Of the 47 potential protein coding genes, 26 were homologous to “hypothetical proteins” found in other fungi, but no known function was assigned. The remaining 12 candidate transcripts (labelled “unkRNAs”) had no known homology among expressed transcripts found in other organisms, ranged in size from 287 nucleotides (nt) to 3,001 nt in length, and had G:C content ranging from 48% (slightly above the norm for *C. auris*) to as low as 36.9%. Differential gene expression analysis was carried out (as above) and nine of these candidate novel genes were found to be significantly differentially expressed (abs(FC) > 2, FDR p(x) < 0.05) in treated versus untreated conditions (Supplemental Data 4). This suggests that these transcripts may be encoded by genes with an as yet undetermined role in the adaptive stress response to antifungal compounds.

### Deep whole transcriptome profiling of PDR and MDR *C. auris* post-exposure to 5-FC

The whole genome sequencing results suggested perturbations in genes associated with nucleotide biosynthesis, mRNA processing and nuclear export of mRNA from the nucleus. Disruption of these genes would be expected to have strong impacts on the entire transcriptome, not solely in response to drug treatment but potentially also in the absence of drug exposure or stress. To investigate further, we carried out whole transcriptome sequencing of two strains (20-34, Day 4 and 20-32, Day 69) both exposed to 5-FC (12). Transcriptional profiles of each strain under both conditions was carried out with biological triplicates.

Over 1 billion reads were generated, with an average of 85.1 million reads generated per replicate (Table 2). An average of 99.39% of reads mapped to the reference genome (Table 2), enabling deep characterisation of the transcriptional response to 5-FC. Expression patterns for replicates appeared similar within each strain (Figure 2A). Principal Component Analysis (PCA) revealed replicates for each condition clustered together, indicating a high level of data correlation, but with a strong separation for each of the four conditions, suggesting genetic differences between 20-34 and 20-32 had strong effects on basal transcriptomes even in the absence of drug treatment (Figure 2B). Gene expression levels for all genes are shown in Supplementary Data 2.

**Figure 2:**
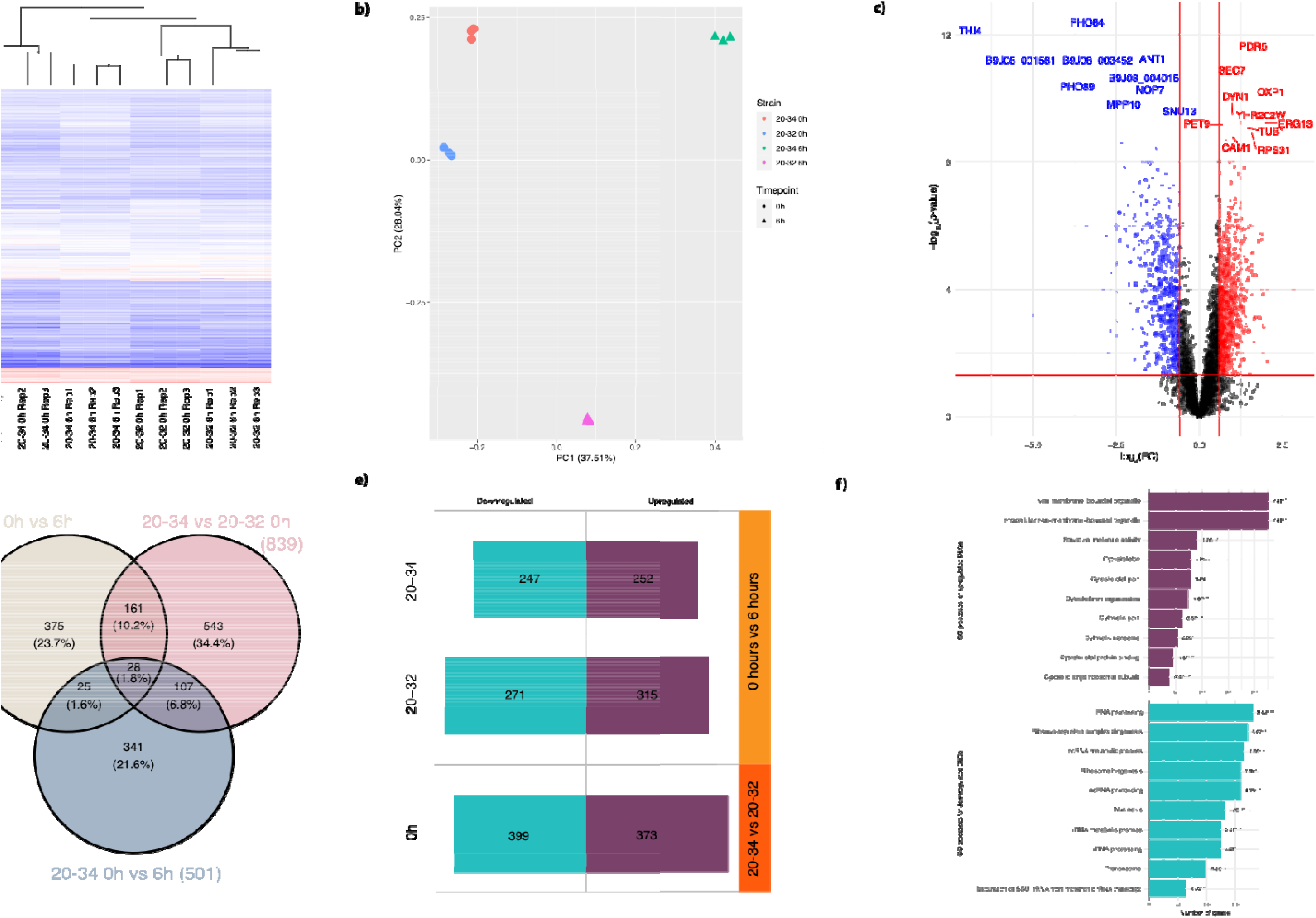
Transcriptome analysis of MDR *C. auris* strain (20–34) and PDR strain (20–32) pre-and 6 hours post-exposure to 5FC. Gene expression profiles of all *C. auris* strains were determined using RNAseq of three biological replicates. A) Heatmap of hierarchical clustering and FPKM values in all strains. Log_2_ FPKM+1 values are color-coded according to the legend at the top left. B) Principal Component Analysis (PCA) of RNAseq read counts from three biological replicates per strain. Clustering of replicates per strain indicates a high level of data correlation, yet with strong separation on PC1 and PC2 axes. C) Adjusted *p*-value versus log fold change (log2(FC)) for all differentially expressed genes. Significantly differentiated genes are denoted as either ‘upregulated’ in red, or ‘downregulated’ in blue. The vertical lines denote the log_2_FC thresholds (-0.6 and 0.6), and the horizontal line denotes the adjusted *p*-value threshold (0.05). The top 10 up and down-regulated genes are labelled. D) Venn diagrams depicting the overlap of differentially expressed genes between the pan-resistant strain (20–34) and susceptible strain (20–32) at pre and 6 hours post-exposure to 5FC. E) Total changes in the *C. auris* transcriptome. Numbers inside the turquoise and purple indicate the number of downregulated and upregulated, respectively, differentially expressed genes in each pairwise comparison. F) Overrepresented GO processes in the *C. auris* transcriptome. The top panel (upregulated DEGs; purple) and bottom panel (downregulated DEGs; turquoise) show the number of genes associated with the top overrepresented GO terms, and associated significant adjusted *p*-value (FDR).

**Table 2:**
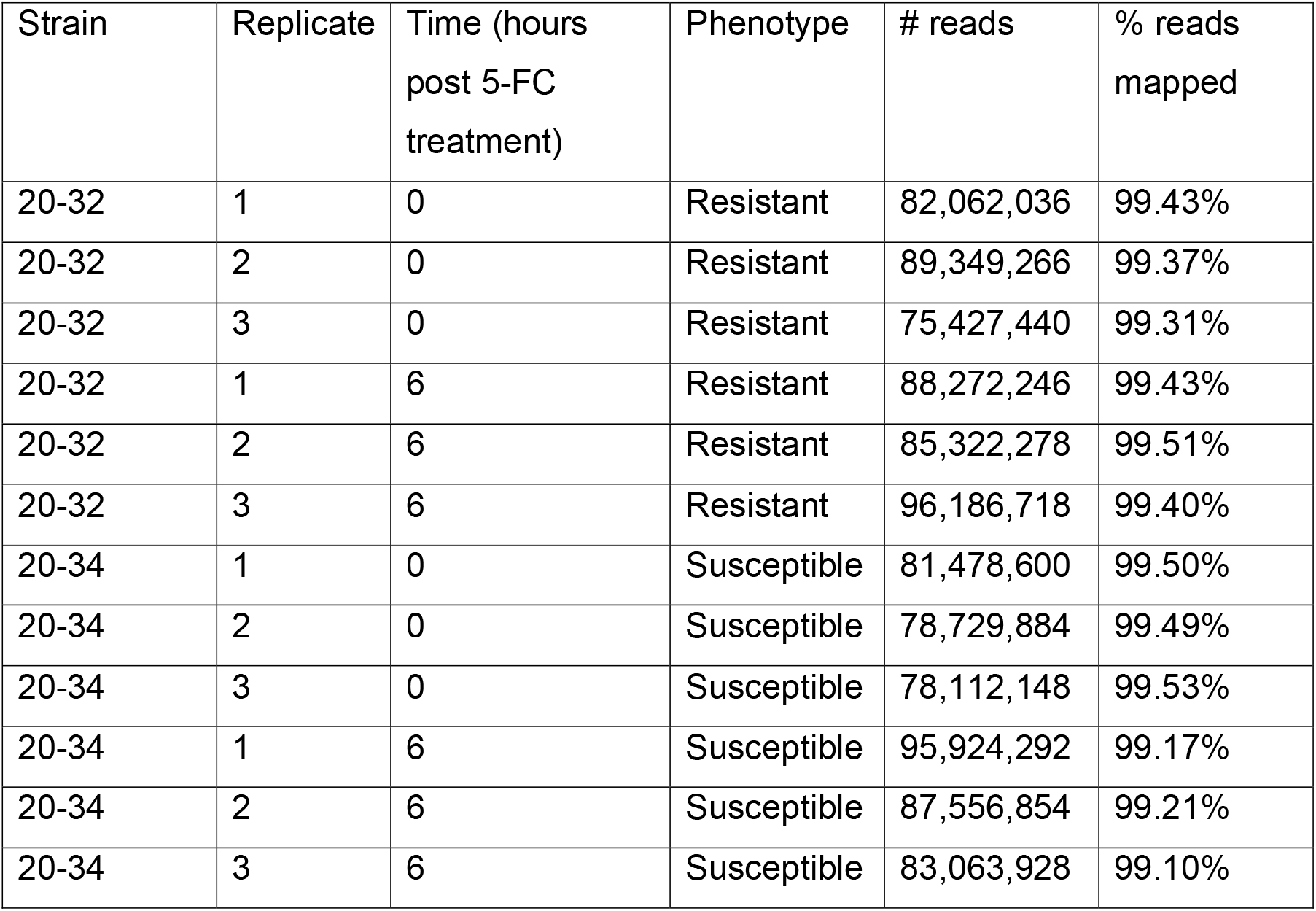
Details of RNAseq alignments for each *C. auris* strain and replicate.

In the principal component analysis (PCA) 20-34 at 6 hours was separated from both 20-32 and 20-34 at 0 hours by the first PC, explaining roughly 38% of the variation (Figure 2B). In contrast, 20-32 at 6 hours was separated by the other strains and timepoints by the second PC, explaining another 28% of the variation (Figure 2B). We considered genes with a *p*-value < 0.05 as differentially expressed genes (DEG), with log_2_FC > 0.6 as upregulated and log2FC < −0.6 as downregulated. In total, 839 genes were significantly differentially expressed, with 373 upregulated and 399 downregulated (Table 3, Figure 2C, Supplementary Data 3). Further analysis revealed that prior to drug treatment with 5-FC, MDR *C. auris* 20-34 (Day 4) and PDR *C. auris* 20-32 (Day 69) showed remarkably different transcriptome profiles from one another; a total of 839 genes, representing over 14% of the entire transcriptome, were found to be differentially regulated (Supplementary Data 3). Of these, 543 DEGs were unique to this response, and not found to be differentially expressed in other conditions (Figure 2D). 93 genes were also extremely dysregulated, displaying log fold changes <-3 or >3. Twenty-eight genes were found to be significantly differentially expressed in all three comparisons: B9J08_003855, a DNA repair protein and B9J08_004693 (*CAC2*) (Figure 2D). Orthologs of B9J08_003855 contain a BRCT domain, which is involved in cell cycle checkpoint functions responsive to DNA damage. *CAC2* is a subunit of chromatin assembly factor 1 (CAF-1), along with rlf2 and Msi1p, which are important for the deactivation of the DNA damage checkpoint after DNA repair.

**Table 3:** Proportions of up and downregulated differentially expressed *C. auris* genes. % values rounded to 1 decimal place.

| *C. auris*<br>comparison | Total | Up (%) | Neutral (%) | Down (%) |
| --- | --- | --- | --- | --- |
| 20-34 0 vs 6h | 501 | 252 (50.3) | 2 (0.4) | 247 (49.3) |
| 20-32 0 vs 6h | 589 | 315 (53.5) | 3 (0.5) | 271 (46.0) |
| 20-34 vs 20-32<br>0h | 839 | 373 (44.5) | 67 (8.0) | 399 (47.5) |
| 20-34 vs 20-32<br>6h | 271 | 115 (42.4) | 1 (0.4) | 155 (57.2) |

GSEA analysis of the 839 DEGs revealed enriched GO terms for oxidoreductase activity (52 genes, corrected *p*-value 3.42^e-06^) for downregulated genes and sulphur amino acid metabolic process (11 genes, corrected *p*-value 2.39^e-02^) for upregulated genes. Of the 93 highly dysregulated genes, the upregulated genes were enriched for uracil phosphoribosyltransferase activity (*p*-value 4.27^e-03^, adjusted *p*-value not significant). Downregulated dysregulated genes were enriched for metal ion transport however, specifically copper (2 genes, adjusted *p*-value 4.08^e-02^). Of these genes, *CTR2* and *FRE2*, both of which have roles in copper ion transport, were extremely downregulated (log_2_(FC) of −3.577 and −3.322 respectively).

Following 5-FC treatment, both strains displayed a large degree of transcriptome-wide remodelling after 6 hours, with 20-34 having 501 DEGs and 20-32 having 589 DEGs compared to their pre-exposed states, with roughly equal numbers up and downregulated (Table 3). However, only 53 DEGs were shared between 20-34 and 20-32 following treatment with 5-FC; only 25 of these were unique to these two strains following 5-FC, the remaining 28 were also found in both strains prior to treatment. Of the 375 DEGs specific to the response of 20-32 to 5-FC, GSEA revealed nucleotide and purine binding dominating the list, and drug metabolism KEGG pathway enrichment. Broad transcriptomic differences were observed in both strains before and after drug treatment with 5-FC (Figure 2E).

GSEA of all DEGs revealed enrichment of RNA processing events in downregulated DEGs, such as alternative splicing and RNA editing, and enrichment for structural processes including the cytoskeleton and non-membrane bounded organelles in upregulated genes (Figure 2F).

### Response to 5-FC by known resistance and biofilm-associated genes

Next, we investigated the transcriptional response to genes previously identified as associated with resistance to 5-FC. Following exposure to 5-FC, the transcriptome of MDR *C. auris* 20-34 (Day 4) included enrichment for KEGG drug metabolism cytochrome P450 pathway. Notably, *FUR1*, a target of 5-FC, was found to be differentially expressed and upregulated as expected in 20-34 post treatment with 5-FC. However, *FUR1* was not differentially expressed in PDR *C. auris* 20-32, both before and after 5-FC treatment, confirming that the 131-nucleotide deletion effectively eliminated its expression in this strain. In addition, multiple hallmark genes previously shown to be associated with resistance were present in both strains, including genes involved in the ergosterol pathway and efflux pump biosynthesis (Figure 3A). The ergosterol biosynthesis gene *ERG11* (B9J08_001448) was considered a DEG in both strains but in 20-32 it was upregulated 2x, and in 20-34 was downregulated 2x fold following 5-FC treatment. Notably, a Lys143Arg (K143R) mutation was present in both strains; this mutation has been shown to be strongly associated with fluconazole resistance (43).

**Figure 3:**
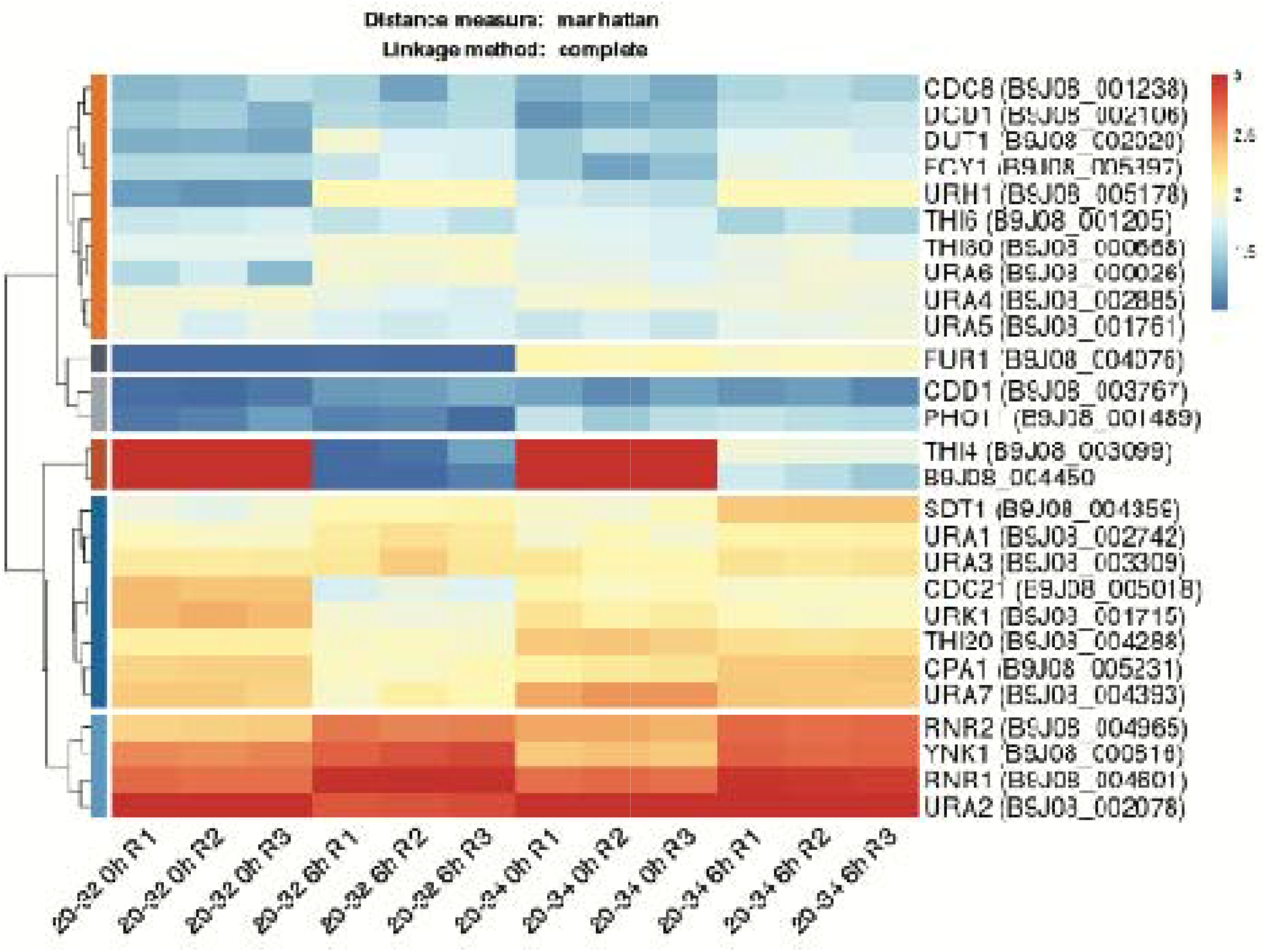
Pyrimidine metabolism. CPM expression values were transformed log10(x), with ∈(-inf, 0] to NAs. Data was clipped to a range of [1,3] (blue lines). Agglomerative hierarchical clustering was performed using R’s standard dist and hclust functions, with a dissimilarity matrix between sample pairs calculated using a Manhattan distance measure. Clusters are successively built and merged together, with the distance determined using a complete linkage method.

**Figure 4:**
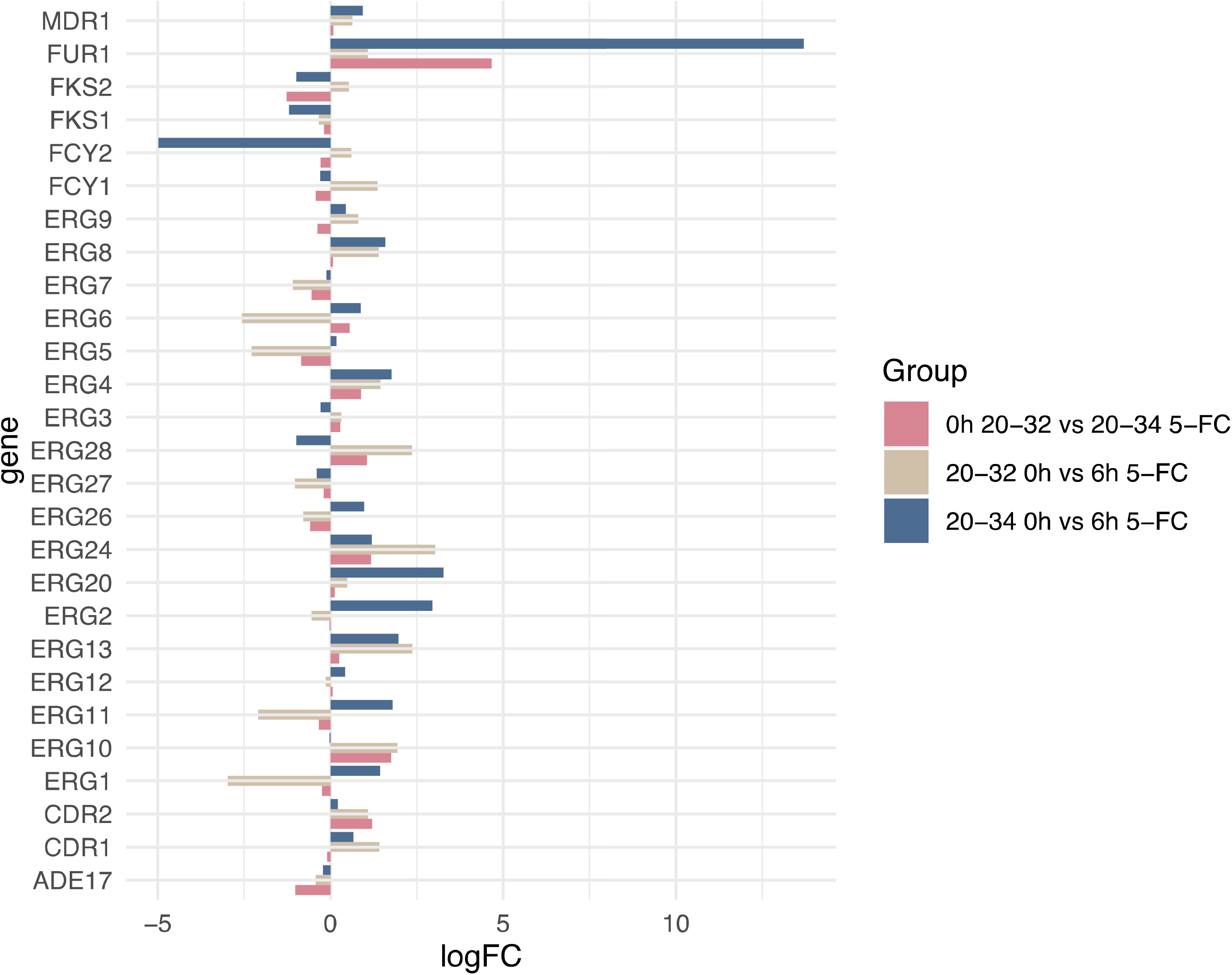
Expression profiles for selected resistance genes in all three comparisons. Profiles depict the log10 fold change.

Following exposure to 5-FC, the transcriptome of the Day 4 strain (20–34) included 12 DEGs associated with pyrimidine-containing compound metabolic processes (GO:0072527, FDR *p*-value 0.03, 12/27 genes). This same GO term was not considered enriched in the Day 69 strain 20-32 (FDR *p*-value 0.50, 9/27 genes), but did share in the 9 DEGs genes with 20-32 (Figure 3A). Notably, *FUR1* was found to have near zero expression in 20-32, both before and after 5-FC treatment (Figure 3), confirming that the 131-nt deletion effectively eliminated its expression in this strain. We also observed differential expression of genes associated with biofilm formation (Table 4), which have also been implied to be associated with differences in drug susceptibility (26).

**Table 4:** DEGs associated with biofilm formation in *C. auris* strains (20-32 and 20-34) following drug treatment with 5-FC.

| Gene ID | Gene name | Regulation |
| --- | --- | --- |
| B9J08_001196 | *CSA1* | Down |
| B9J08_000458 | *CEK1* | Down |
| B9J08_000928 | *AQY1* | Down |
| B9J08_003614 | *ADH2* | Down |
| B9J08_000003 | *Zrt1* | Up |
| B9J08_004334 | *PHO2* | Up |
| B9J08_001624 | *ILS1* | Up |
| B9J08_001939 | *RIX7* | Up |
| B9J08_002043 | *ARO1* | Up |
| B9J08_005492 | *QDR1* | Down |

### Difference in growth and phenotype profile of pan-drug resistant *C. auris*

We found differences in *in vitro* growth of two *C. auris* strains. *Candida auris* 20-32 consistently outgrew *C. auris* 20-34 on both RPMI and SAB media at 30 °C and 37 °C after 48 hours, and this was significant at both 30 °C and 37 °C for SAB and for RPMI at 37 °C (Figure 5). Phenotypic profiling on various carbon and nitrogen sources was compared for *C. auris* 20-32 and 20-34. A total of 190 carbon and 95 nitrogen sources were included in the test panels. Robust growth was estimated at absorbance cut-off of 0.5. Nitrogen sources were preferred by both *C. auris* 20-32 (47 substrates) and 20-34 (42 substrates). Relatively less vigorous growth was seen on a variety of carbon sources with only 16 of 190 carbon sources supporting growth above cut-off for *C. auris* 20-32 and 11 carbon sources for *C. auris* 20-34. Among carbon sources key to central metabolism, sorbitol, mannitol, and N-acetyl-D-glucosamine) impacted upper glycolysis while acetate/acetic acid and citrate/citric acid were key elements in the tricarboxylic acid cycle (TCA)(44) (Figure 5).

**Figure 5:**
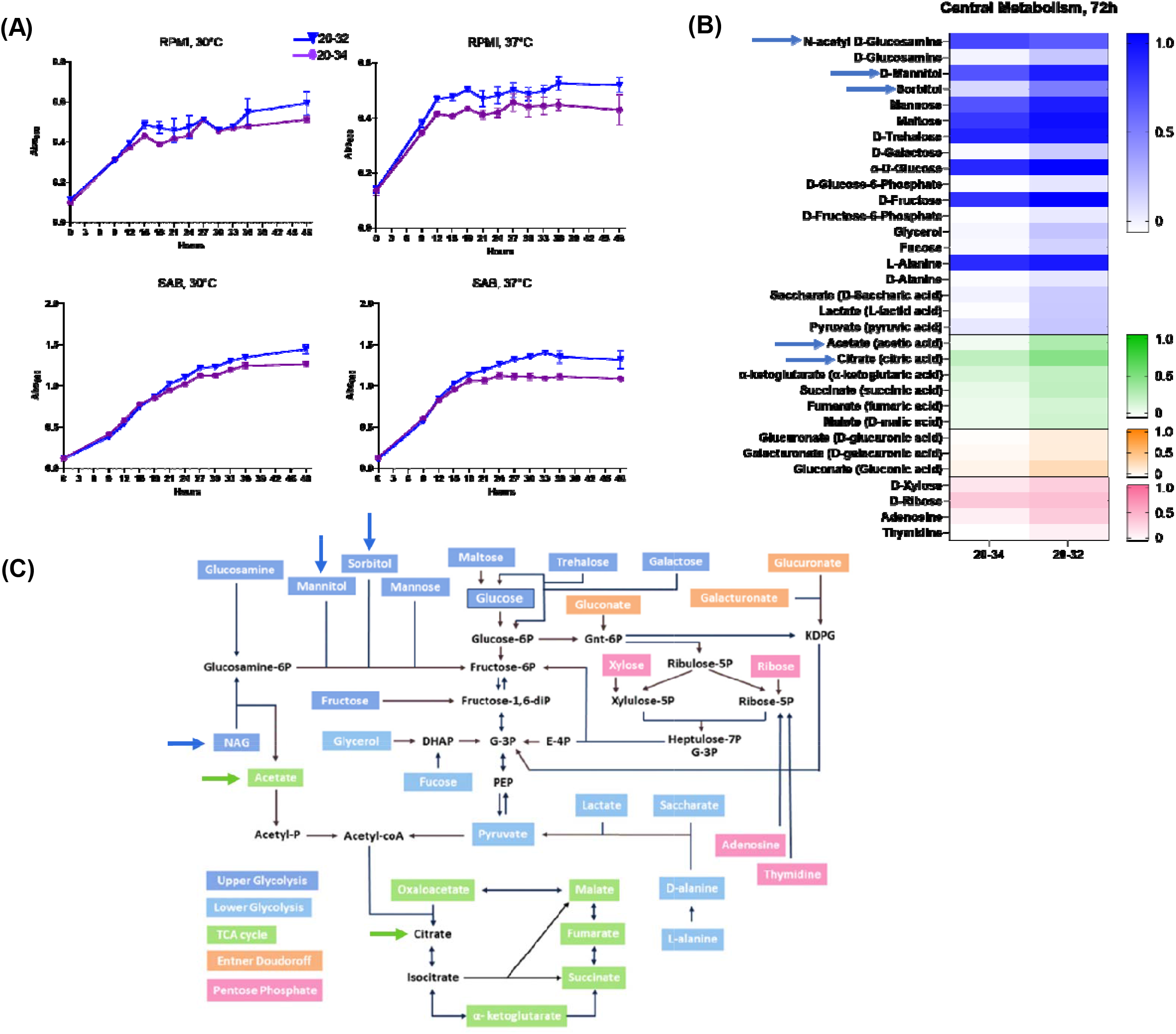
Fitness and metabolic profiles of *C. auris*. (A) Growth curves at 30 °C and 37 °C show comparable growth in nutrient-deficient RPMI and nutrient-rich SAB media. (B) Phenotypic profiling on various carbon (190 sources) and nitrogen (95)sources. Nitrogen sources were preferred by PDR *C. auris* 20-32 (47 substrates) and MDR 20-34 (42 substrates); carbon source growth was less vigorous for *C. auris* 20-32 (16 substrates) and *C. auris* 20-34 (11 substrates). (C). Among carbon sources key to central metabolism, sorbitol, mannitol, and N-acetyl-D-glucosamine) impacted upper glycolysis while acetate/acetic acid and citrate/citric acid were key elements in the tricarboxylic acid cycle (adapted from Tong et al. (44).

### Computational modelling suggests alternate routes for uracil monophosphate generation

The *C. auris* UPRT tetramer reactant state model with uracil and PRPP suggests how 5-fluorouracil might be processed by *C. auris* UPRT (figure 6). The binding site for 5-fluorouracil is likely to be identical to that of uracil in each monomer structure. Once bound in this location next to a PRPP molecule, it would then be catalytically converted, in the same fashion as uracil, to 5-fluorouracil monophosphate. The N-terminal 33 amino acids (aa) in each *C. auris* UPRT monomer (shown in yellow, Figure 6) participate extensively in interactions with neighbouring monomers. While their absence could modify substrate binding allosterically, it should more directly affect tetramer formation. The UPRT mutant with the 33 aa N-terminal deletion therefore may have diminished activity. This suggests some alternate route in *C. auris* to generate uracil monophosphate that, in contrast to UPRT, does not accommodate 5-fluorouracil as a substrate.

**Figure 6:**
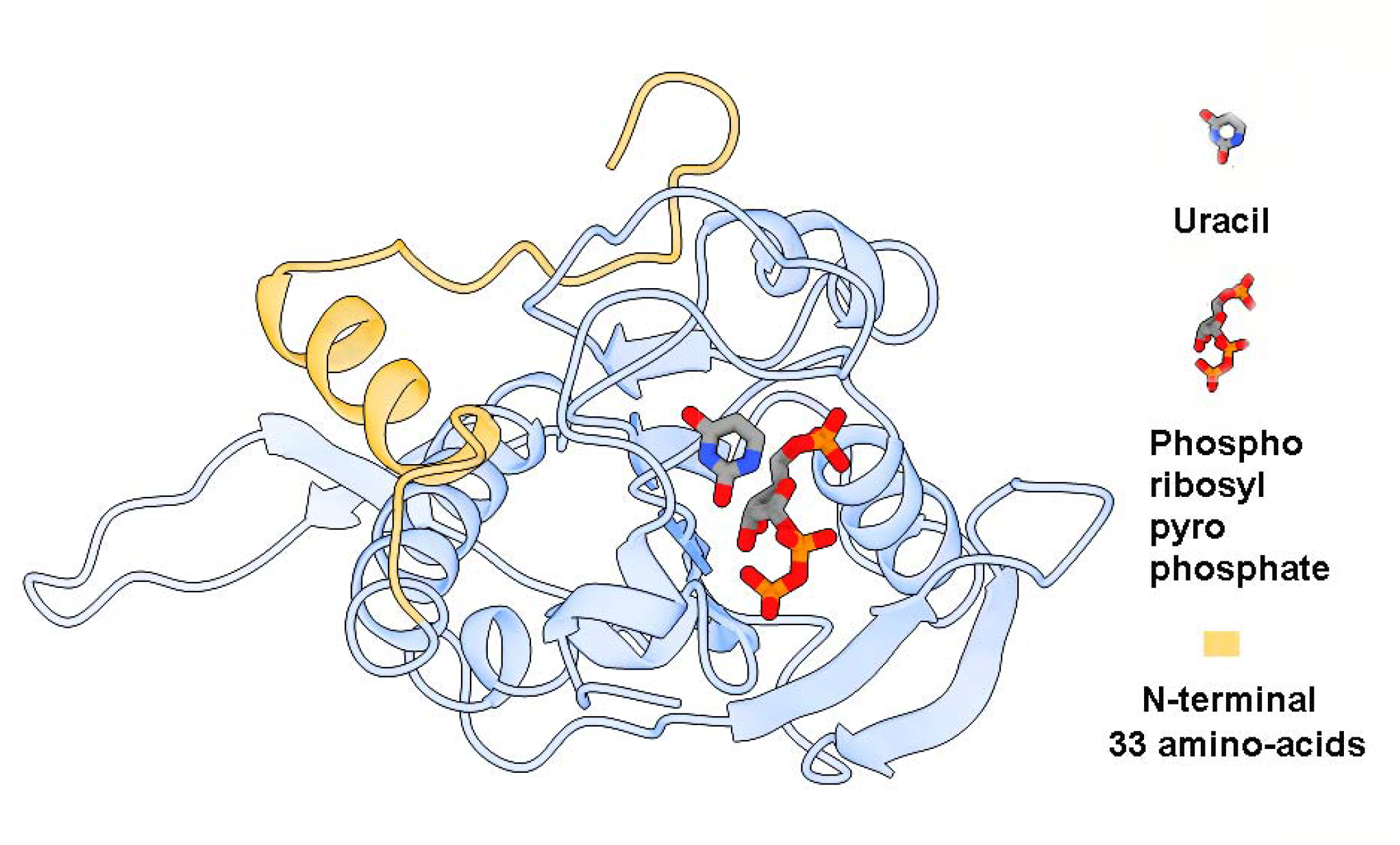
A reactant state model for the *C. auris* uracil phosphoribosyl transferase (UPRT) monomer. The two substrates, uracil, and phosphoribosyl pyrophosphate, are shown in stick format in a *C. auris* UPRT monomer. The 33 amino acids at the N-terminus, shown in yellow, are not directly interacting with the active site, but do participate in interactions with neighboring monomers in the tetramer state.

## Discussion

*Candida auris* represents a severe public health threat globally, causing severe nosocomial infections. Our research, utilizing comparative genomic, RNAseq, and phenotypic analyses on two strains, each representing different levels of antifungal drug resistance, has unveiled a unique set of metabolic adaptations. These findings are unprecedented and have led to the identification of unique genomic, transcriptomic, and phenomic signatures. Upon data curation, we have discovered both similarities and differences between the MDR and PDR *C. auris* strains before and after treatment with 5-FC. In recent years, there is renewed interest in the use of 5-FC in antifungal combination therapy, but there remains a major concern around the development of 5-FC resistance by fungal pathogens. Therefore, it is not just important, but absolutely crucial that we comprehend the mechanisms governing resistance to 5-FC in this significant and emerging fungal pathogen, *C. auris*.

Previously, *C. auris* 20-32 had displayed a pan-drug resistance profile. In our transcriptomic analysis we confirm *FUR1*, the target of 5-FC, was not differentially expressed both before and after treatment, confirming the impact of the 131-nucleotide deletion. *FUR1* was, however, expressed in *C. auris* 20-34, which did not display a pan-drug resistance profile. This was further confirmed by GO terms enriched for pyrimidine-containing compound metabolic processes in MDR strain 20-34, but not in the PDR strain 20-32. Genomic analysis and subsequent GSEA showed 20-32 significantly enriched for pyrimidine-containing compound salvage pathway and poly(A)+ mRNA export from the nucleus, which suggests selective pressure from the therapeutic regiment pushed the *C. auris* population towards acquiring mutations in these key pathways. *Candida glabrata* strains with partial deletions in *FUR1* have also been demonstrated to confer 5-FC/5FU cross-resistance. It is likely the mutations conferring 5-FC resistance cause inactivation of enzyme activity in the pyrimidine salvage pathway, thus no longer inhibiting *C. auris*, due to no redundancy in the pathway in *FCY1* and *FCY2*.

Here, we found almost 1,000 differentially expressed genes in response to 5-FC treatment. Previous studies have identified a number of DEGs in relation to drug resistance in *C. auris*; for example, a study by Muñoz *et al.* identified 39 and 21 DEGs in susceptible *C. auris* strains exposed to amphotericin B and voriconazole, respectively, and 14 DEGs were common to both responses (23). Additionally, 38 and 18 were repressed by amphotericin B and voriconazole, respectively, and 9 were repressed in response to both drugs (23). Of these, the DEGs common to both responses were also found to be differentially expressed in response to 5-FC. However, this represents a small number of DEGs in response to drug. Zhou *et al.* identified almost 2000 DEGs when comparing drug resistant to drug susceptible *C. auris* (25). Of these, 1913 were also found to be differentially expressed in response to 5-FC, and all the common DEGs from Muñoz *et al.* were also found to be common, suggesting a common regulatory network in response to drug pressure in *C. auris* (23). The strong separation of the four conditions observed in the PCA suggests that genetic differences between 20-34 and 20-32 pre and post-exposure to 5-FC had strong effects on the transcriptomes (Fig 2A and B). Previously, distinct transcriptomic profiles have been observed in echinocandin-resistant *C. auris* strains, suggesting a core signature gene set linked to resistance to this particular drug (27). The smaller number of DEGS when comparing 20-34 to 20-32 after exposure to 5-FC may also represent a conserved gene regulatory network for the response to this drug, which may be expanded upon in other conditions. Further research into the transcriptional regulation of the genes in this pathway, in addition to the changes occurring at the membrane proteome level, are required to understand the full spectrum of the mechanisms of 5-FC resistance.

Comparison of *C. auris* 20-32 with 20-34 revealed relatively similar fitness in the pandrug-resistant phenotype as observed in nutrient-enriched and -deficient media at 30°C and 37°C. Our observations are consistent with an earlier report on the absence of any fitness cost for acquisition of fluconazole resistance in *C. auris* (26), and contrary to two recent reports on drug-resistance-associated fitness defect in *C. auris* (45, 46). *C. auris* has previously demonstrated significant increases in the utilisation of specific carbon sources (47), and we observed both 20-32 and 20-34 preferred nitrogen sources over carbon. It is difficult to reconcile these findings as variable methods were used for growth curve measurements in the published studies. Broadly, the correlation between fitness cost and drug-resistance remains equivocal in pathogenic *Candida* species (48–51). The emerging evidence suggests that unlike bacterial drug-resistance mechanisms, pathogenic fungi might undergo metabolic adaptions with increased fitness in drug-resistant phenotypes (52, 53). While metabolic differences have not yet been associated with drug-resistance in *C. auris*, species-specific responses have been demonstrated on various carbon and nitrogen sources (54, 55). These unique metabolic features could not only contribute to its emergence as a pathogen, but also its spread, as fungal resistance does not equate to a fitness compromise.

Whilst micafungin has been recommended as the first-line treatment for *C. auris* in adults, it is clear we need to rise to the challenge of increasing antifungal drug resistance profiles in this pathogen. Several new antifungal agents have been developed and are currently undergoing clinical trials, such as ibrexafungerp, rezafungin, and fosmanogepix. Urgent research needs to be conducted to discern whether resistance mechanisms to existing antifungal drugs confer resistance to these novel antifungal drugs; this will enable us to stay afoot of potential resistance development to these novel antifungal drugs to ensure successful treatment of *C. auris* infections.

## Acknowledgements

We thank technical expertise of Matt Shudt, Applied Genomic Technologies Core, and Jonathan Adams, Media and Glassware Core, Wadsworth Center, New York State Department of Health, Albany, NY, USA. Supported in part by a Centres for Disease Control and Prevention-Antibiotic Resistance Lab Network grant (NU50CK000423) and a grant (1R21AI156573-01A1) from the National Institutes of Health. JR is supported by a Welcome Trust fellowship (219551/Z/19/Z). The transcriptomic analysis was performed using the High-Performance Computing facility and Research Data Store at Imperial College London (DOI: 10.14469/hpc/2232).

---

**Supplementary Figure 1:** Mutations present in all three Day 69 strains, but not found in the Day 4 strain A) Predicted deletion of 131 bp (PIS52459.1:c-38_101del) in *FUR1* B) Frameshift mutation resulting in Ser527fs near the terminus of the protein encoded by B9J08_001846 is present in all three Day 69 strains.

**Supplementary Data**

Supplementary Data 1 (Excel format): details of 299 variants unique to Day 69 strains when compared to Day 4, of which 82 were common to all three Day 69 strains.

Supplementary Data 2 (Excel format): raw FPKM and coverage for each transcript (and associated gene) for each condition and replicate.

Supplementary Data 3 (Excel format): differentially expressed genes for *C. auris* strains 20-34 and 20-32 prior to drug treatment with 5-FC, with details on regulation.

