## Supplementary figures and images for "What makes *Candida auris* pan-drug resistant? Integrative insights from genomic, transcriptomic, and phenomic analysis of clinical strains resistant to all four major classes of antifungal drugs"

### Supplementary Figure

#### Slide 1
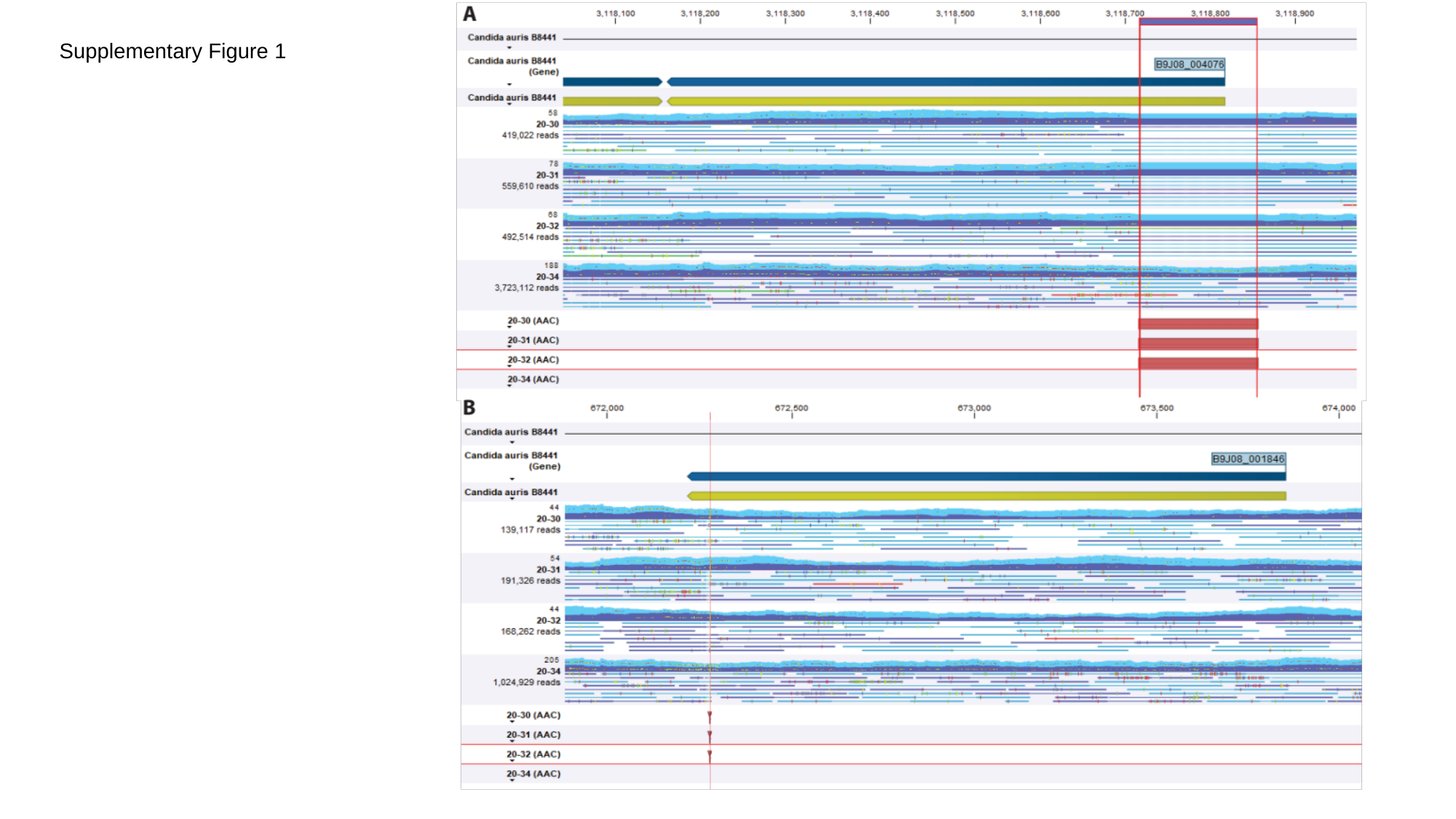

Supplementary Figure 1
